# Demographic fluctuation of community-acquired antibiotic-resistant *Staphylococcus* aureus lineages: potential role of flimsy antibiotic exposure

**DOI:** 10.1101/206078

**Authors:** Claude-Alexandre Gustave, Anne Tristan, Patricia Martins-Simoes, Marc Stegger, Yvonne Benito, Paal Skytt Andersen, Michèle Bes, Philippe Glaser, Frédéric Laurent, Thierry Wirth, François Vandenesch

**Author notes:** Corresponding author: François Vandenesch, CIRI (International Center for Infectiology Research), Inserm U1111/CNRS UMR5308, Team “Staphylococcal pathogenesis”, Domaine de la BUIRE, 8 Rue Guillaume Paradin, 69372 Lyon cedex 08, FRANCE.

## Abstract

Community-acquired (CA) -as opposed to hospital acquired- methicillin-resistant *Staphylococcus* aureus (MRSA) lineages arose worldwide during the 1990s. To determine which factors, including selective antibiotic pressure, govern the expansion of two major lineages of CA-MRSA, namely “USA300” in Northern America and the “European ST80” in North Africa, Europe and the Middle East, we explored virulence factor expression, and fitness levels with or without antibiotics. The sampled strains were collected in a temporal window representing various steps of the epidemics, reflecting predicted effective population size as inferred from whole genome analysis. In addition to slight variations in virulence factor expression and biofilm production that might influence the ecological niches of theses lineages, competitive fitness experiments revealed that the biological cost of resistance to methicillin, fusidic-acid and fluoroquinolone is totally reversed in the presence of trace amount of antibiotics. Our results suggest that low-level antibiotics exposure in human and animal environments contributed to the expansion of both European-ST80 and USA300 lineages in community setting. This surge was likely driven by antibiotic (ab)use promoting the accumulation of antibiotics as environmental pollutants. The current results provide a novel link between effective population size increase of a pathogen and a selective advantage conferred by antibiotic resistance.

## INTRODUCTION

*Staphylococcus aureus* remains one of the most common causative agents of both nosocomial and community-acquired infections. It colonizes asymptomatically about one third of the human population and may cause infections with outcomes ranging from mild to life-threatening (Lowy, 1998). Until the mid-1990’s, methicillin-resistant *S. aureus* (MRSA) infections were reported almost exclusively from hospital settings and most hospital-associated MRSA (HA-MRSA) diseases resulted from a limited number of successful clones (Thurlow *et al.*, 2012). These HA-MRSA, which remained confined to healthcare settings, were exposed to a high antibiotic pressure among patients with frequent immunity impairment and/or invasive devices such as urinary/vascular catheters or mechanical ventilation (Chavez and Decker, 2008). Therefore, HA-MRSA were likely under strong positive selection within these healthcare-associated niches where the acquisition of resistance to multiple antibiotic families provided them with a major competitive advantage despite their impaired fitness. However, in the beginning of 2000’s, MRSA infections began to be reported in healthy individuals without known risk factors or apparent connections to healthcare institutions (Chambers, 2001), (Vandenesch *et al.*, 2003). These community-acquired (CA)-MRSA strains had genetic backgrounds distinct from the traditional HA-MRSA strains with specific lineages predominating in different continents such as the Sequence Type 8 (ST8) SCCmecIVa (standing for <u>s</u>taphylococcal <u>c</u>assette <u>c</u>hromosome encoding <u>m</u>ethicillin resistance gene of type IVa) pulsotype USA300 in the USA (abbreviated to “USA300” below), the ST80 SCCmecIV in Europe, North Africa and the Middle East (hereinafter referred to as “EU-ST80“), and the ST30 SCCmecIV in Oceania (Mediavilla *et al.*, 2012). Some genetic features of these CA-MRSA were postulated to be major determinants of their selective advantages against HA-MRSA in community settings (David and Daum, 2010). Fitness impairment associated with the SSC*mec* mobile element is a well-described example; large SCC*mec* elements shared by HA-MRSA induced a stronger fitness decrease compared to small SCC*mec* of CA-MRSA (Ma *et al.*, 2002), the latter being therefore promoted under lightened antibiotic pressure outside of healthcare settings. Successful community spread of CA-MRSA has also been allegedly associated with ecological factors such as modifications of colonisation niches. This was illustrated by the hypothesis of a deleterious impact of anti-pneumococcal vaccines on nasal microbiota facilitating CA-MRSA colonization (Regev-Yochay *et al.*, 2006). Finally, the observation that CA-MRSA had apparently increased virulence for human (Li *et al.*, 2010) notably in skin infection, suggested that higher bacterial load associated with increased severity of cutaneous infections could promote dissemination between humans.

Regarding the population dynamics of CA-MRSA, recent phylogenetic studies, conducted on the USA300 (Glaser *et al.*, 2016) and the EU-ST80 (Stegger *et al.*, 2014) lineages, proposed two Bayesian evolutionary models inferring their population size through time among hundreds of isolates sampled from 1980’s to 2000’s. Those phylogenetic analyses strongly suggested that in the transition from an MSSA lineage to a successful CA-MRSA clone, the USA300 lineage first became resistant to multiple antibiotics, acquired the arginine catabolic mobile element (ACME) which encodes factors promoting skin colonization and infection (Thurlow *et al.*, 2013), and subsequently acquired resistance to fluoroquinolones (Planet *et al.*, 2015). These two steps were associated with two successive phases of sharp demographic expansion of what is known as the USA300 North-American (NA) lineage as opposed to the Latin-American Variant (LV) which does not harbor ACME (Glaser *et al.*, 2016). A similar study, performed on the EU-ST80 epidemic CA-MRSA lineage, depicted a clone derived from a Panton-Valentine (PVL)-positive methicillin-susceptible *S. aureus* (MSSA) ancestor from sub-Saharan Africa that dramatically expanded in the early 1990’s once out of West Africa, upon acquisition of the SCC*mec* element, the plasmid-encoded fusidic-acid resistance (*fusB*) and four canonical SNPs including a non-synonymous mutation in the accessory gene regulator C (*agrC*) (Stegger *et al.*, 2014), a major virulence regulator in *S. aureus* (Reynolds and Wigneshweraraj, 2011). However, for both the USA300 and the EU-ST80 lineage it remains to be demonstrated that the identified genetic events, which correlate with the demographic expansion, are causally related with population size variations. In order to answer these questions, we explored fitness and virulence factor expression of strains selected at various evolutionary and temporal stages of the predicted population size inferred through Bayesian coalescence models.

## MATERIALS & METHODS

### Strain selection

Infection-related strain selection among the CA-MRSA clones “USA300” and “EU-ST80” was determined by two previously published phylogenic studies (Glaser *et al.*, 2016), (Stegger *et al.*, 2014). All isolates were stored at −20°C at the National Reference Center for Staphylococci (NRCS - HCL, Lyon), on cryobeads. Prior whole genome sequencing and Bayesian analysis of all strains enabled their assignment to an evolution phase of these clones; thus isolates of the two lineages were selected at various temporal steps of their inferred population dynamics as follows: for the CA-MRSA USA300 lineage, ten clinical strains plus one reference strain were included (Fig. 1a & Table 1): (i) two strains corresponding to the most recent common ancestor of the USA300 clone, lacking the ACME sequence (Basal USA300 1 & 2), (ii) four strains from the early expansion phase characterized by ACME and SCC*mec* acquisition (Derived USA300 1, 2, 3 & 4), (iii) four strains from the most recent evolutionary phase subsequent to fluoroquinolones resistance acquisition (Derived USA300 5, 6, 7 & 8). For the EU-ST80 lineage, eleven clinical isolates plus one reference strain were selected (Fig. 1b & Table 1): (i) five strains from the basal clade with a high genetic proximity with their hypothetical common MSSA ancestor from Sub-Saharan Western Africa (Basal MSSA 1, 2, 3, 4 & 5), (ii) two MRSA strains from the derived clade isolated from a patient from Maghreb (Derived MRSA 1 & 2), (iii) two MRSA strains from the derived clade isolated on patients from Europe (Derived MRSA 3 & 4), (iv) two MRSA strains from the derived clade and associated with the stabilization/decline phase of the lineage (Derived MRSA 5 & 6); The competitive strain pairs are summarized in Table S1. Reference strain for the USA300 and EU-ST80 lineages were SF8300-LUG2295 and HT20020209-LUG1799 respectively (Table 1).

**Figure 1:**
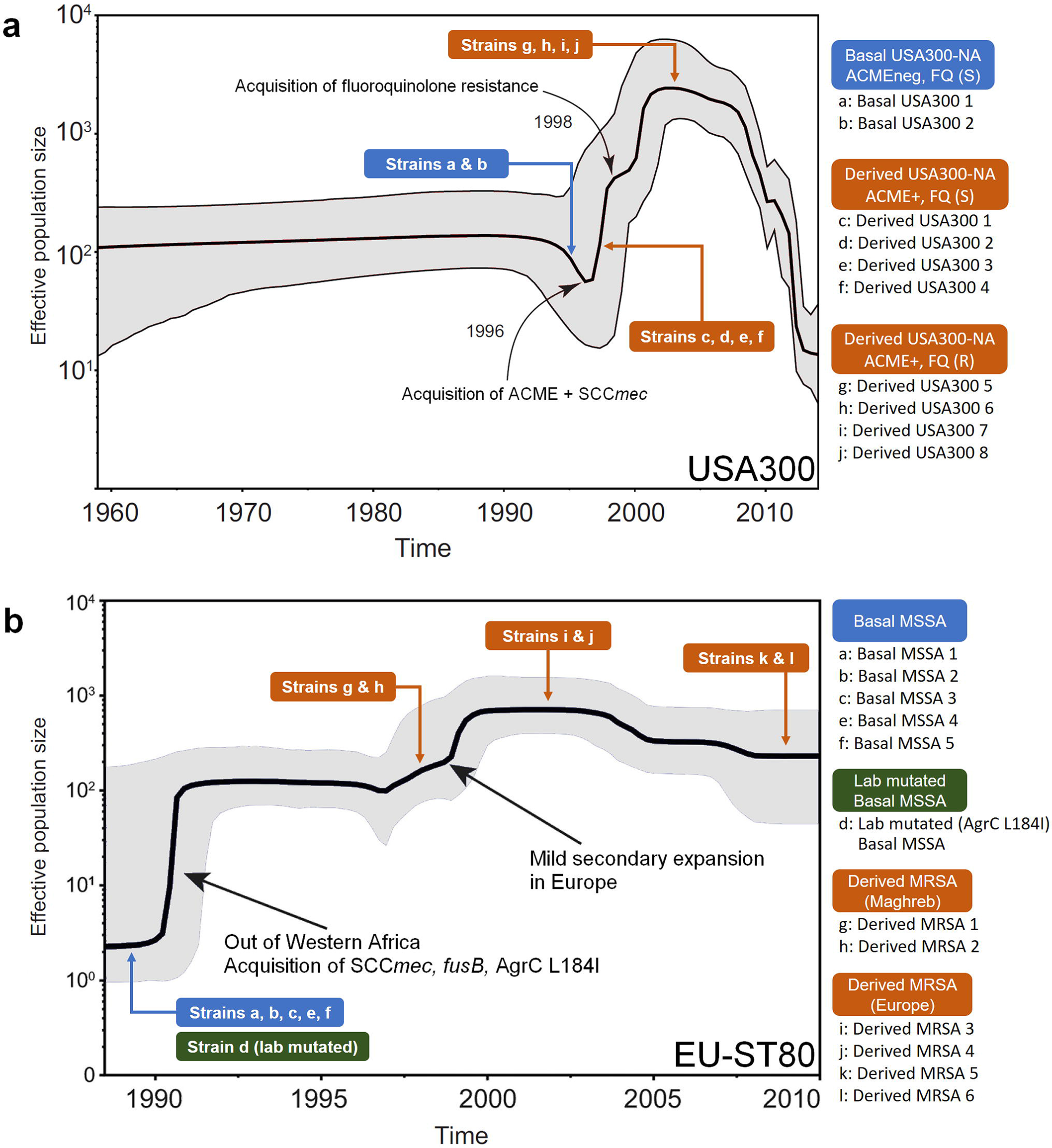
Bayesian demography of USA300 and EU-ST80 lineages. Bayesian skyline plot indicating population size changes in the USA300 **(a)** and EU-ST80 **(b)** lineages over time with a relaxed molecular clock. The shaded area represents the 95% confidence interval. Strain selection and their designation are indicated by colored thumbnails. Adapted from Glaser *et al.* (2016) and Stegger *et al.* 2014.

**Table 1:** Relevant characteristics of strains. Strains list with their ID’s (as used in the manuscript and figures), NRCS Number (as used in reference publications), relevant characteristics, and reference publications.

| STRAIN DESIGNATION | NRCS number | RELEVANT CHARACTERISTICS | REFERENCES |
| --- | --- | --- | --- |
| **<u>USA300 strains</u>** |  |  |  |
| USA300 ref strain | SF8300-LUG2295 | *mecA*, ACME, [R = Oxa, E, C, T, Cip, Mup] | Diep *et al.*, 2008 |
| Basal USA300 1 | ST2003-0343 | *mecA*, [R = Oxa] | This study |
| Basal USA300 2 | ST2005-0026 | *mecA*, [R = Oxa] | This study |
| Derived USA300 1 | ST2012-0558 | *mecA*, ACME, [R = Oxa] | Glaser *et al.*, 2016 |
| Derived USA300 2 | ST2012-1514 | *mecA*, ACME, [R = Oxa] | Glaser *et al.*, 2016 |
| Derived USA300 3 | ST2013-0343 | *mecA*, ACME, [R = Oxa,K,E] | Glaser *et al.*, 2016 |
| Derived USA300 4 | ST2011-2484 | *mecA*, ACME, [R = Oxa,K,E] | Glaser *et al.*, 2016 |
| Derived USA300 5 | ST2011-1414 | *mecA*, ACME, [R = Oxa,K,E,O] | Glaser *et al.*, 2016 |
| Derived USA300 6 | ST2013-0068 | *mecA*, ACME, [R = Oxa,K,E,O] | Glaser *et al.*, 2016 |
| Derived USA300 7 | ST2013-1284 | *mecA*, ACME, [R = Oxa,K,E,O, T] | Glaser *et al.*, 2016 |
| Derived USA300 8 | ST2013-1763 | *mecA*, ACME, [R = Oxa,K,E,O, T] | Glaser *et al.*, 2016 |
| **<u>EU-ST80 strains</u>** |  |  |  |
| EU-ST80 ref strain | HT20020209-LUG1799 | *mecA*, AgrC L184, [R = Oxa, K, T, F] | Perret *et al.*, 2012 |
| Basal MSSA 1 | HT2002-0042 | AgrC L184 | Stegger *et al.*, 2014 |
| Basal MSSA 2 | HT2006-0859 | AgrC L184 | Stegger *et al.*, 2014 |
| Basal MSSA 3 | HT2003-0006 | AgrC L184, [R = T] | Stegger *et al.*, 2014 |
| Lab Mutated Basal MSSA | LUG2417 | **AgrC L184I**, [R = T] | This study |
| Basal MSSA 4 | HT2004-1302 | AgrC L184, [R = T] | Stegger *et al.*, 2014 |
| Basal MSSA 5 | HT2005-0374 | AgrC L184, [R = T] | Stegger *et al.*, 2014 |
| Derived MRSA 1 | ST2007-1277 | *mecA*, **AgrC L184I**, [R = Oxa] | Stegger *et al.*, 2014 |
| Derived MRSA 2 | ST2007-1273 | *mecA*, **AgrC L184I**, [R = Oxa] | Stegger *et al.*, 2014 |
| Derived MRSA 3 | ST2005-0508 | *mecA*, **AgrC L184I**, [R = Oxa,K,F] | Stegger *et al.*, 2014 |
| Derived MRSA 4 | ST2009-0942 | *mecA*, **AgrC L184I**, [R = Oxa,K,F,T] | Stegger *et al.*, 2014 |
| Derived MRSA 5 | ST2007-0258 | *mecA*, **AgrC L184I**, [R = Oxa,K,F,T,E,C] | Stegger *et al.*, 2014 |
| Derived MRSA 6 | ST2007-1047 | *mecA*, **AgrC L184I**, [R = Oxa,K,F,T,E,C] | Stegger *et al.*, 2014 |

C: Clindamycin - Cip: Ciprofloxacin - E: Erythromycin - K: Kanamycin - NRCS: National Reference Center for Staphylococci - O: Ofloxacin - Oxa: Oxacillin - R: resistant - T: Tetracycline - F: Fusidic acid – I: Isoleucine – L: Leucine - NRCS: National Reference Center for Staphylococci - Oxa: Oxacillin - R: resistant

### Construction of agrC mutant

The *agrC* locus of one basal MSSA of the ST80 lineage (Basal MSSA 3) (Fig. 1b & Table 1) was mutated by allelic replacement to confer the sequence carried by isolates of the derived clade (isoleucine instead of leucine at position 184). This mutation was obtained by using pMAD (Arnaud *et al.*, 2004). Two *agrC* DNA fragments flanking the *agrC* target region were amplified from a wild type strain using agrC2912/agrC555 and agrC544/agrC4238 primers respectively (Table S2). DNA fragments were then blunt-ended by *Sca*I and *Pvu*II restriction enzymes, before being ligated and amplified using external primers agrC2912/agrC4238. The resulting DNA fragment corresponding to an *agrC* encoding sequence for the mutated amino-acid 1184 was restricted by *Xho*I and *Pvu*II and cloned in pMAD linearized by *Sal*I and *Sma*I. The resulting plasmid, pLUG1166, was electroporated into RN4220, and then into Basal MSSA 3. Transformants were grown at non-permissive temperature (42°C), to select for cells with chromosome-integrated plasmid by homologous recombination. Successful double crossover mutants were subsequently selected on X-gal agar plates after single colony culture at 30°C for 10 generations. PCR amplifications and sequencing were used to confirm the mutation of *agrC* in the resulting strain LUG2417 designated “Lab mutated basal MSSA” (Table 1).

### RNA extraction from S. aureus

Brain-Heart Infusion broth (BHI) was inoculated with an overnight culture to an initial OD_600nm_ of 0.05 and grown up on aerated Erlenmeyer flask to the end of exponential phase (6h) at 37°C under agitation (200 rpm). One milliliter of bacterial suspension was harvested and concentration adjusted to an OD_600nm_ = 1.0. Bacteria were washed in 10 mM Tris buffer and treated with lysostaphin and β-mercaptoethanol. RNAs were extracted with the RNeasy Plus Mini Kit^®^ (Qiagen), quantified by spectrophotometry and stored at −80°C. This process was repeated on three different days for biological replicates.

### RNA quantification by real-time PCR

A random-primers based reverse transcription of 1μg of RNA was performed with the A3500 Reverse Transcription System Kit (Promega), followed by quantitative real-time PCR on cDNA using the FastStart Essential DNA Green Master kit (Roche) and the LightCycler^®^ Nano (Roche). As previously described (Li *et al.*, 2010), we targeted five virulence genes (*RNAIII*, *lukS-PV*, *hla*, *hlgC*, *psmα*) and the housekeeping gene *gyrB* for normalization. Gene expression levels were compared between our clinical isolates and against the reference strains (SF8300-LUG2295 for the USA300 clone and HT20020209-LUG1799 for the EU-ST80 clone); levels were expressed as n-fold differences relative to reference strains or an isolate from another evolutionary phase. These qRT-PCR were performed as technical triplicates (three RNA quantification per RNA sample), on RNA obtained from three biological replicates (three independent cultures and extractions per strain).

### Biofilm production assay

Each isolate was incubated overnight on blood agar (Columbia) at 35°C under ambient air. Three colonies were transferred into 9mL of BHI and incubated with agitation (200 rpm) overnight at 35°C under ambient air. Bacterial suspensions were then placed in a 96-well plate and incubated at 35°C under ambient air for, 24 and 48h respectively. Biofilm production was assessed by spectrophotometry after well drying and crystal violet fixation. *S. aureus* laboratory strains SH1000 was added as a positive control (Horsburgh *et al.*, 2002), (O’Neill, 2010), *S. epidermidis* ATCC12228 (Zhang *et al.*, 2003) and *S. carnosus* TM300 (Rosenstein *et al.*, 2009) as negative controls of adhesion. Biofilm experiments were performed as technical replicates (three wells per strain) and biological replicates (three independent plate series).

### MIC determination

In order to adjust their concentrations in selective media and broth used for sub-inhibitory antibiotic pressure, MIC of second line antibiotics were measured by E-tests on Mueller-Hinton agar according to EUCAST specifications.

### Crude doubling time

Isolates growth curves were determined from BHI cultures incubated in 96-well plates for 24 hours at 37°C with continuous optical density monitoring at 600nm (Tecan Infinite^®^ 200 PRO). Each strain was inoculated in three independent wells (technical replicate), and the experiment was repeated on three different days (biological replicate). Doubling times were calculated by graphical method with the Log-transformed optical density data of the exponential growth phase.

### Competitive fitness

Each strain to be tested in a competitive pair was adjusted to an OD_600nm_ = 1, then 3 mL of a 1/100 dilution in BHI of each strain was mixed in a glass tube. For some experiments ofloxacine, ceftriaxone, or fusidic acid were added at final concentrations corresponding to 1/4 to 1/100 of the susceptible strain’s MIC. Tubes were incubated at 35°C in aerobic atmosphere under agitation (200 rpm) for 22 +/− 2h, and 50 μL were transferred daily for 21 days to a fresh tube containing 3 mL of BHI. The proportion of each strain in the competitive mix was monitored daily with differential colony count based on selective agar inoculated with a calibrated amount of competitive mix (Spiral System^®^ - Interscience) followed by aerobic incubation for 24h at 37°. For MSSA vs MRSA couples, we used the ChromAgar^®^ medium (i2A, France) allowing for growth of both strains (total count) and the ChromlD-MRSA^®^ medium (BioMérieux, France) for MRSA colony count. For the USA300 clone, where all isolates were MRSA, we used second line antibiotics resistance for strain discrimination. Therefore, differential colony counts were performed with simultaneous inoculation of a brain-heart agar (BHA) and a BHA with ofloxacin (2 μg/mL, i.e. x5 above sensitive strain MIC, x6 below resistant strain MIC). Similarly, for MRSA vs MRSA pairs belonging to the EU-ST80 lineage, a combination of BHA and BHA with tetracycline (1 μg/mL, x8 above sensitive strain MIC, x8 below resistant strain MIC) was used. Strain quantifications calculated from colony counts on selective agar were confirmed by quantitative PCR targeting discriminant genes (*mecA* for MSSA versus MRSA, *grlA* for fluoroquinolones sensitive versus fluoroquinolones resistant, *tetK* for tetracycline sensitive versus tetracycline resistant or *arcA*-ACME for ACME negative versus ACME positive strains) carried by one of the strains in the competitive pair. This approach was used to rule out a growth inhibition bias on selective medium. This was also the only strain quantification method usable for the EU-ST80 Basal MSSA 1 in competition with its *agrC* derivative obtained by allelic replacement. Strain proportions were determined with a L184I-specific set of primers (Table S2). All the PCRs were performed at days 0, 7, 14 and 21. Continuous competitive cultures were performed on three independent series (biological triplicates), each colony count or qPCR was performed on three technical triplicates. For all strains pairs tested, one of the strains was eventually reduced to a trace level, so no statistical test was required for strains proportions comparison.

## RESULTS

### Growth rate along the phylogeny

Previous studies showed that CA-MRSA grew significantly faster than HA-MRSA, a property that may be a prerequisite for CA-MRSA, in the absence of antibiotic pressure, to achieve successful colonization of humans by outcompeting the numerous bacterial species in the human environment outside the hospital setting (Okuma *et al.*, 2002). We thus tested whether growth rate assessed by doubling-time varied between isolates of USA300 and EU-ST80 CA-MRSA lineages selected at various temporal steps of their Bayesian demography (Fig. 1 and Table 1). The experiment performed on ten USA300 isolates pointed out the impact of ACME acquisition on doubling time shortening (Derived USA300 1 or 2 versus Basal USA300 1 or 2, Mann-Whitney test, *P* = 0.029) (Fig. 2a). Within the derived clade corresponding to the epidemic phase, crude fitness appeared to be fading as we observed significant increase of doubling time along the phylogeny as shown by intraclade comparisons (Mann-Whitney test, *P* = 0.029 for all comparisons) (Fig. 2a). Each doubling time rise appeared to be related to a new acquisition of antibiotics resistance, namely aminoglycosides and macrolides, then fluoroquinolones, followed by tetracyclines (Fig. 2a and Table 1). Among the twelve strains belonging to the EU-ST80 CA-MRSA lineage that were tested, the shortest doubling times were observed for the “basal clade” isolates (interclade comparison, Mann-Whitney test, *P* = 0.029) (Fig. 2b). Within each clade (basal and derived), we observed a decreasing crude fitness along the phylogeny as shown by increasing of doubling times (intraclade comparisons, Mann-Whitney test, *P* = 0.029 for all comparisons) (Fig. 2b). Like for USA300 strains, antibiotics resistance appeared to be a major determining factor of doubling time lengthening as shown by interclade comparison (Basal MSSA vs Derived MRSA, Mann-Whitney test, *P* = 0.029), and by intraclade comparisons revealing longer doubling times associated with new acquisition of antibiotics resistance, namely tetracyclines within the basal clade of MSSA strains; whereas in the derived clade, fitness impairments resulted from the consecutive acquisition of resistance to beta-lactams, fusidic acid, aminoglycosides, and finally tetracyclines and macrolides for the most recent isolates (Mann-Whitney test, *P* = 0.029 for all comparisons) (Fig. 2b and Table 1). At this stage, as epidemic strains (from derived clades) displayed the longest doubling times, we concluded that crude *in vitro* fitness did not explain the evolutionary dynamic of the two lineages. We therefore investigated other features related to host interaction and antibiotic pressure that could explain the demography of both lineages.

**Figure 2:**
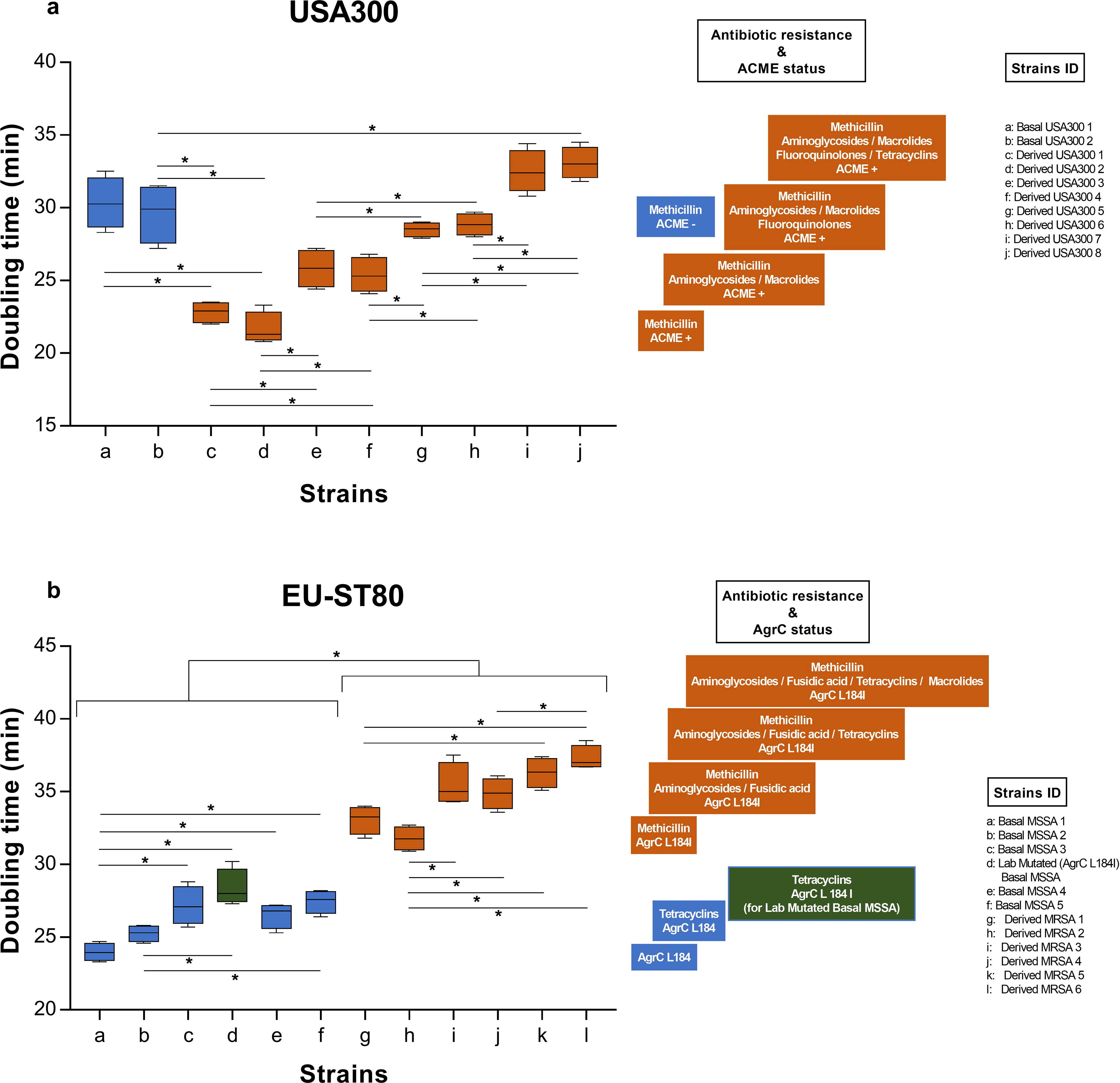
Doubling times of USA300 and EU-ST80 strains. USA300 **(a)** or EU-ST80 **(b)** isolates were cultured in BHI incubated on 96-wells plates for 24 hours at 37°C with continuous optical density monitoring at 600nm (Tecan Infinite^®^ 200 PRO). Doubling times were calculated by graphical method after Log transformation of data from the exponential growth phase. The color codes for each strain correspond to those in Fig. 1. (*: *P* = 0.029). Experiments were performed on three independent series (biological replicates), and optical densities were measured on three wells for each strain (technical replicates).

### Expression of core-genome encoded virulence factors along the lineages’ evolutionary history

Previous studies revealed that overexpression of core-genome encoded virulence factors was a common feature of CA-MRSA, a characteristic that has been proposed to contribute to the expansion of these lineages (Li *et al.*, 2010). We therefore tested whether variations in the expression of core genome encoded virulence factors along the Bayesian demographic models could be observed (Fig. 1). To this end, RT-PCRs targeting virulence factors of the core-(α-toxin, PSMα, γ-toxin) and accessory-genome (*LukSF-PV*), as well as the major regulator (*agr*-RNAIII), were performed after *in vitro* post-exponential growth as previously described (Li *et al.*, 2010). Among the USA300 CA-MRSA isolates, despite an outlier strain with no measurable expression of *hla*, no major variations were detected in expression levels of the targeted virulence factors (above the accepted 2-fold level generally considered as a minimum biological relevant variation in RT-PCR approaches) between the strains representing the various steps of the demography (Fig. 3a). This lack of significant differences was observed by either using an ancestral strain (Basal USA300 1) or the reference strain SF8300 as comparators (Fig. 3a). For the EU-ST80 lineage, most of the targeted virulence factors studied showed variation in expression below - or close to - two-fold, between ancestral and derived isolates with the exceptions of i) *psm*α increasing by 3 - 3.5-fold in two isolates from the evolutionary-derived clade (designated derived MRSA 2 and 5), and ii) *luk*SF-PV increasing by a factor of 2.8-fold in one isolate (derived MRSA 3) (Fig. 3b). In addition, expression of RNAIII, the *agr*-related regulatory RNA, was slightly increased among isolates of the derived clade (reaching a 2.1-fold increase for one strain) compared to the basal clade (Fig. 3b).

**Figure 3:**
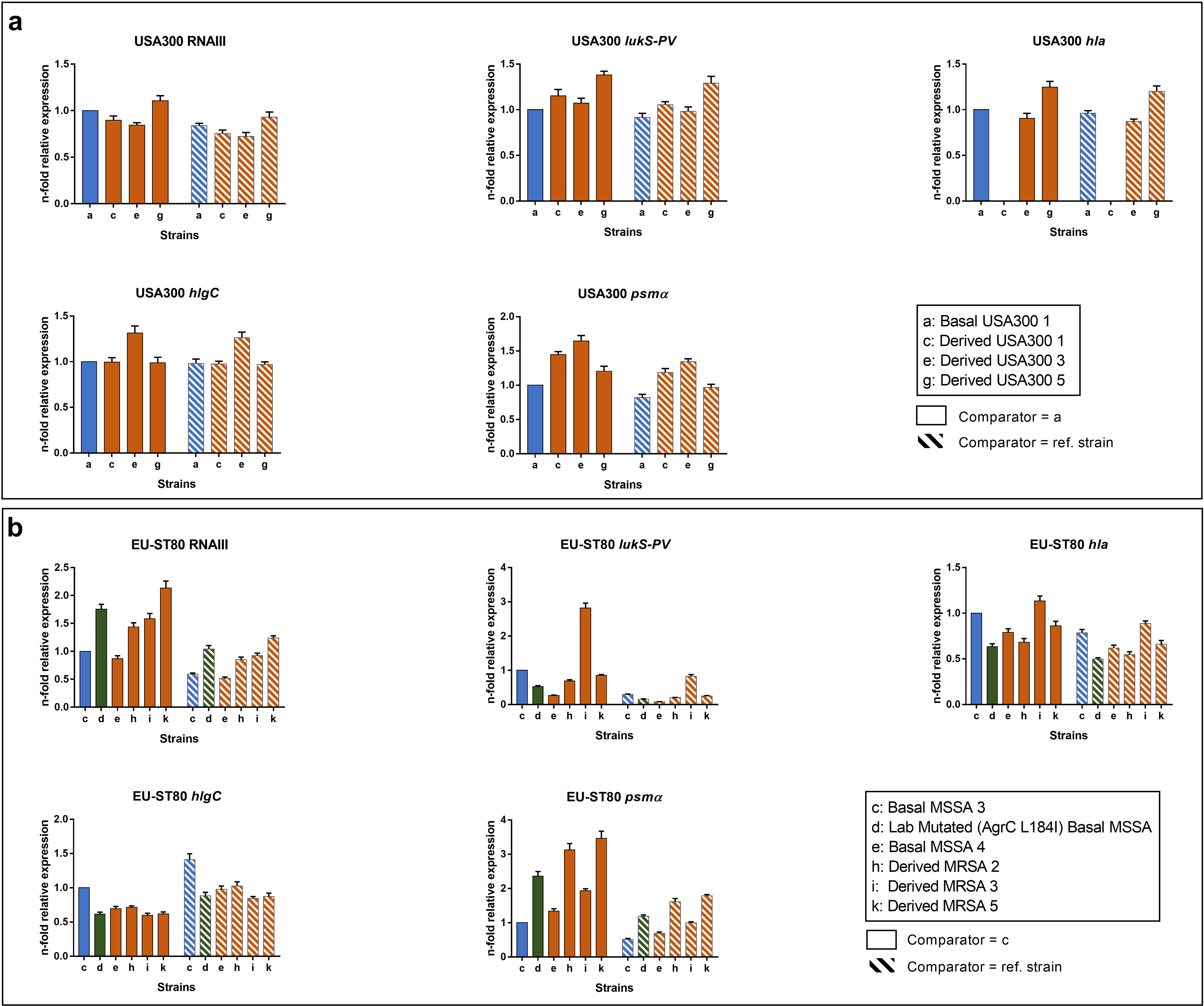
Expression of virulence related genes among USA300 and EU-ST80 strains. Expression of virulence factor- and regulatory-genes were assessed by qRT-PCR among USA300 isolates **(a)** and EU-ST80 isolates **(b)** of various temporal phases of the demographic expansion. Results are expressed as fold change in comparison to the most ancestral strain of the lineage (plain) or to the reference strain of the lineage (striped). Experiments were performed on three independent series (biological replicates), and three RNA quantifications were done for each RNA sample (technical replicates).

All the EU-ST80 isolates from the derived clade harbor an L184I mutation in the extracellular loop of the AgrC receptor (Stegger *et al.*, 2014), a mutation that may have a functional impact on Agr signalling and expression of *agr*-RNAIII. To further investigate this point, an ancestral ST80 (AgrC L184) was engineered by allelic replacement to carry the L184I substitution and was then tested (as “Lab Mutated Basal MSSA“) for quantification of RNAIII and virulence factor expression. Compared to wild type (L184), the mutated (L184I) isogenic derivative (Lab Mutated Basal MSSA) showed a slight enhancement of RNAIII expression, but below the level of 2 (Fig. 3b). This mutation had no significant impact on virulence gene expression, except a mild 2.2-fold increase in *psm*α expression (Fig. 3b).

### Biofilm production

The detection of a slight difference in RNAIII production associated with the *agrC* mutation in the EU-ST80 lineage prompted us to test whether it could translate into differences in biofilm production. After 48 hours of growth, ancestral MSSA strains displayed a higher production of biofilm compared to derived MRSA strains carrying the L184I AgrC substitution (*P* = 0.0002) (Fig. 4). The role of AgrC L184I substitution in this phenotypic difference was confirmed by comparing the ancestral ST80 (AgrC L184) with its isogenic derivative (L184I), the latter showing a significant reduction of biofilm production (*P* < 0.0001). Importantly, the AgrC L184I mutation had no impact on crude fitness since doubling times of Lab Mutated Basal MSSA and its wild type parental strain were similar (Fig. 2b). Therefore, differences observed in biofilm production were not due to growth variations but rather actual differences in biofilm production.

**Figure 4:**
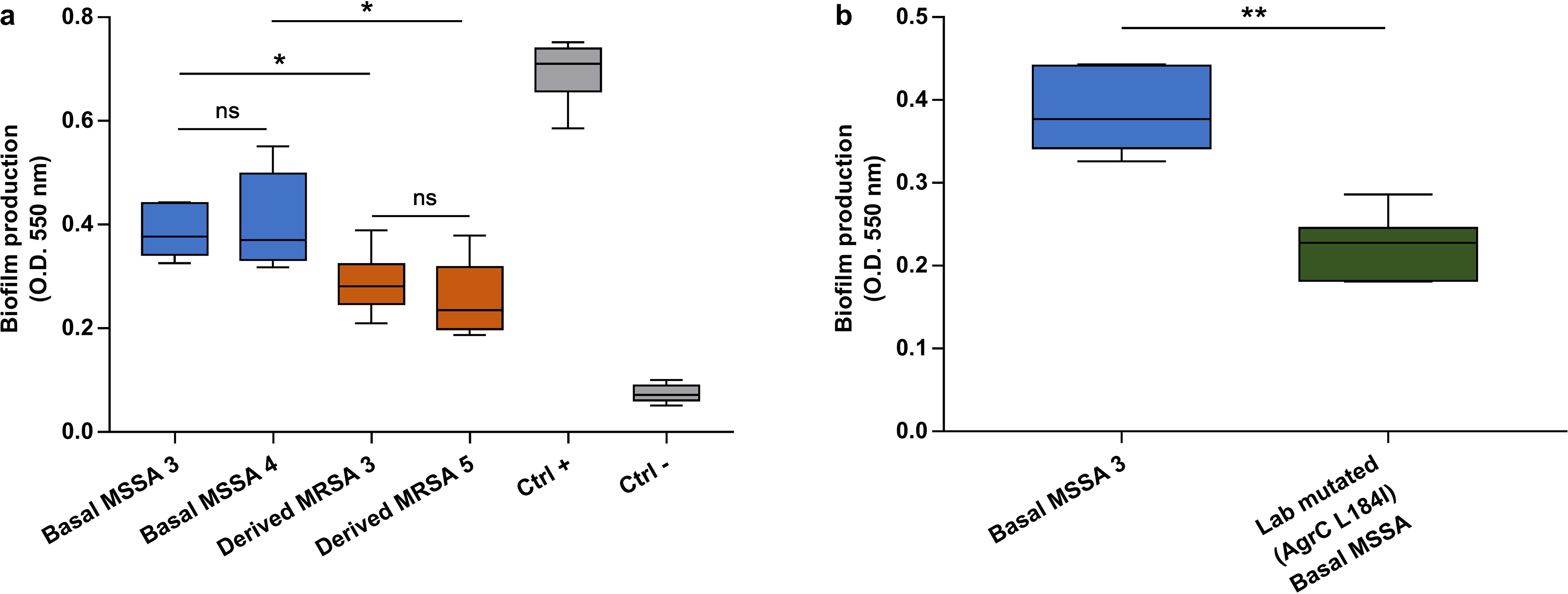
Biofilm production assay for EU-ST80 strains. **(a)** Biofilm production was assessed by crystal violet stain on strains of the EU-ST80 lineage belonging to ancestral clade (Basal MSSA 3 and 4) or derived clade (Derived MRSA 3 and 5), the latter carrying the mecA gene and expressing an AgrC L184I variant. *S. aureus* SH1000 was used as positive control for biofilm production and *S. carnosus* TM300 and *S. epidermidis* ATCC12228 were negative controls, **(b)** Comparison in biofilm production by crystal violet from EU-ST80 Basal MSSA 3 and its isogenic derivative expressing an AgrC L184I variant (Lab Mutated Basal MSSA). The color codes for each strain correspond to those in Fig. 1. (*:*P* = 0.015; **: *P* = 0.002). Experiments were performed on three independent series (biological replicates), and biofilm production was quantified on three wells for each strain (technical replicates).

### Competitive fitness along the phytogeny

Doubling times comparisons used for crude fitness assessment highlighted fitness modulations along the phylogeny that did not match the Bayesian models inferred for both clones (Fig. 1). To better address this issue, we conducted a competitive fitness experiment in more stringent conditions based on continuous co-cultures for 21 days with isolates belonging to each phase of these lineages’ evolution. Competitive strains pairs were designed in order to assess each evolutionary breakpoint identified in their inferred Bayesian phylogenic models (Glaser *et al.*, 2016), (Stegger *et al.*, 2014), (Tables 1 & S1). Moreover, since our assessment of crude doubling times identified the acquisition of antibiotics resistance as a major determinant of fitness alteration, we tested the impact of sub-inhibitory concentrations of antibiotics on competitive fitness of these isolates. Within the USA300 lineage, the acquisition of ACME was associated with an increased fitness: during continuous competitive culture, ACME-positive MRSA strains outcompeted ACME-negative MRSA strains (Fig. 5a). This confirmed the results obtained by crude fitness assessment where shorter doubling-times were obtained with ACME-positive strains compared to ACME-negative ones (Fig. 2a). However, this fitness enhancement was progressively abolished along the phylogeny with the acquisition of fluoroquinolone (FQ) resistance; competitive fitness dropped even below the level observed prior to ACME acquisition: FQ-resistant ACME-positive strain was outcompeted by both FQ-susceptible ACME-positive or-negative strains (Fig. 5b & c). Same results were obtained with a competition between the FQ-resistant strain (Derived USA300 5) and another FQ-susceptible isolate (Derived USA300 4) (data not shown). Altogether these results indicate that ACME enhances fitness but is insufficient to compensate for the fitness cost of FQ resistance.

**Figure 5:**
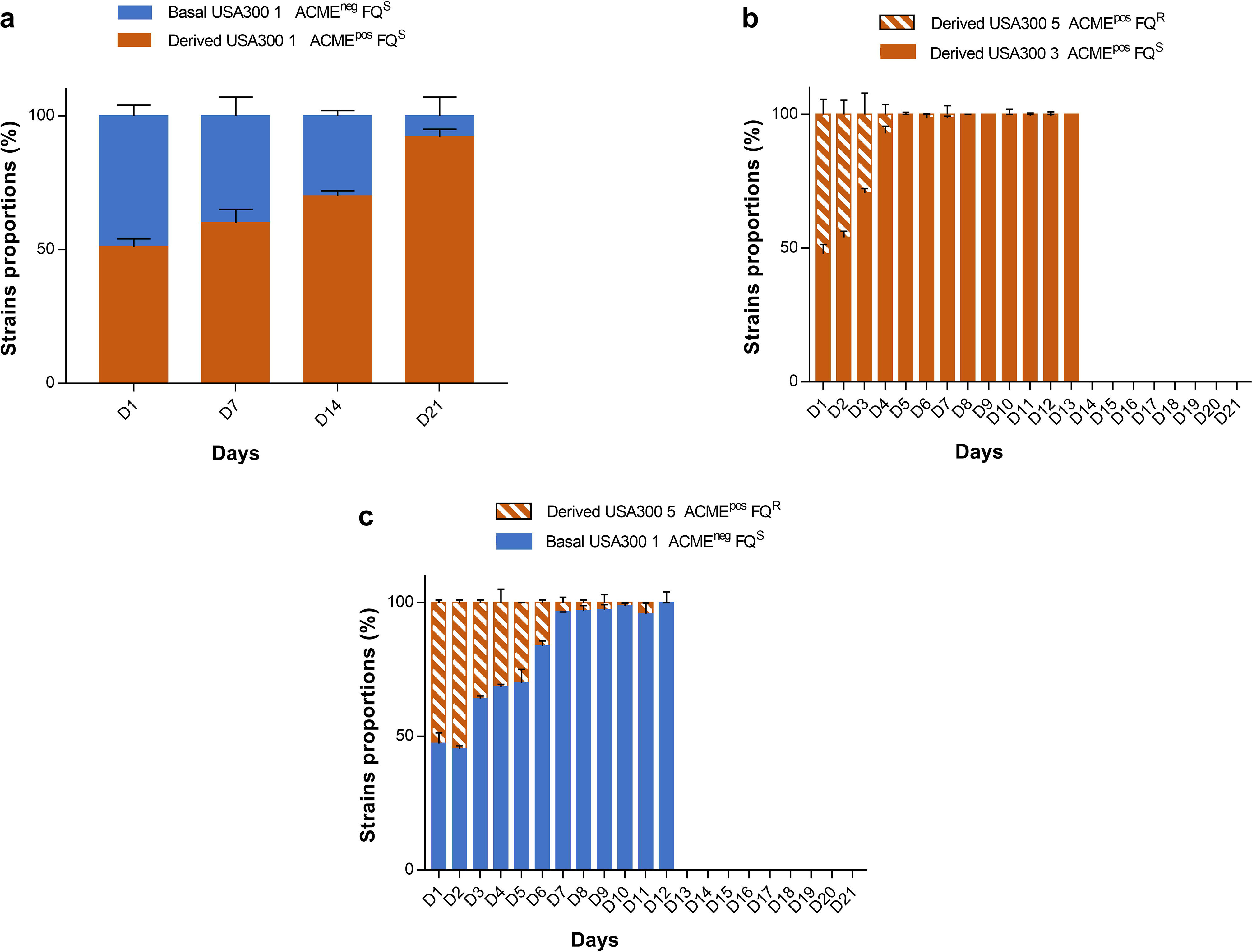
Impact of ACME in competitive fitness of USA300. **(a)** ACM E-negative and -positive strains, both MRSA with no associated antibiotics resistance were co-cultivated for 21 days in BHI with daily subculture in fresh medium. The proportion of each strain was monitored at day 1, 7,14 and 21 with qPCR targeting *arcA*-ACME. **(b)** ACME-positive strains, one Fluoroquinolone (FQ)-susceptible (Derived USA300 3) and one FQ-resistant (Derived USA300 5) were co-cultivated for 21 days in BHI with daily subculture in fresh medium. The proportion of each strain was monitored daily with differential colony count based on selective agar inoculated with a calibrated amount of competitive mix. Same results were obtained with a competition between the FQ-resistant strain (Derived USA300 5) and another FQ-susceptible isolate (Derived USA300 4), data not shown, **(c)** ACME-positive FQ-resistant strain (Derived USA300 5) and ACME-negative strain FQ-susceptible (Basal USA300 1) were co-cultivated for 21 days and assessed as in **(b).** Competitive cultures were performed on three independent series (biological replicates), and each colony count or qPCR was repeated three times (technical replicates).

To assess whether the fitness cost of resistance could be reversed in the presence of trace amounts of antibiotics that could be present in the environment (Okuma *et al.*, 2002), (Gothwal and Shashidhar, 2015), competitive cultures were performed at various sub-inhibitory concentrations of antibiotics. The antibiotics chosen were those those for which resistance acquisition correlate with noticeable variation in effective population size of the lineages (beta-lactams, fusidic acid for EU-ST80, and fluoroquinolones for USA300). Strikingly, even extremely low FQ concentration (1/100 of the FQ-susceptible strain’s MIC, 0.0038 μg/mL) was sufficient to confer a strong selective advantage of FQ-resistant ACME-positive strain toward FQ-susceptible ACME-positive strain (Fig. 6). Similar analyses performed on the EU-ST80 strains also confirmed the results obtained during doubling times assessment suggesting that the major factor ruling the fitness downfall along the phylogeny was not the AgrC L184I but the acquisition of SCC*mec/fusB* and further extended antibiotics resistance. In competitive culture assays, the laboratory engineered *agrC* mutation did not translate into fitness impairment after 21 days of competitive culture with its wild type progenitor (Fig. 7a), whilst competitive culture of the clinical strains confirmed the strong fitness reduction of the derived MRSA isolates compared to the ancestral MSSA in favor of a fitness cost of antibiotics acquisition (the most premature being SCC*mec* and *fus*B) in the absence of antibiotics (Fig. 7b). Similar results were obtained with Basal MSSA 4 versus Derived MRSA 3 (data not shown). The same competition performed in the presence of sub-inhibitory concentration of beta-lactam or fusidic acid totally reversed the result with a strong advantage of the MRSA even at extremely low concentrations (1/100 of MSSA Ceftriaxone MIC, 0.03 μg/mL, and 1/100 of MSSA fusidic acid MIC, 0.0009 μg/mL) (Fig. 7c & d). The same results were obtained with antibiotics concentrations of 1/16 and 1/32 of their MICs and with the couple Basal MSSA 4 versus Derived MRSA 3 (data not shown).

**Figure 6:**
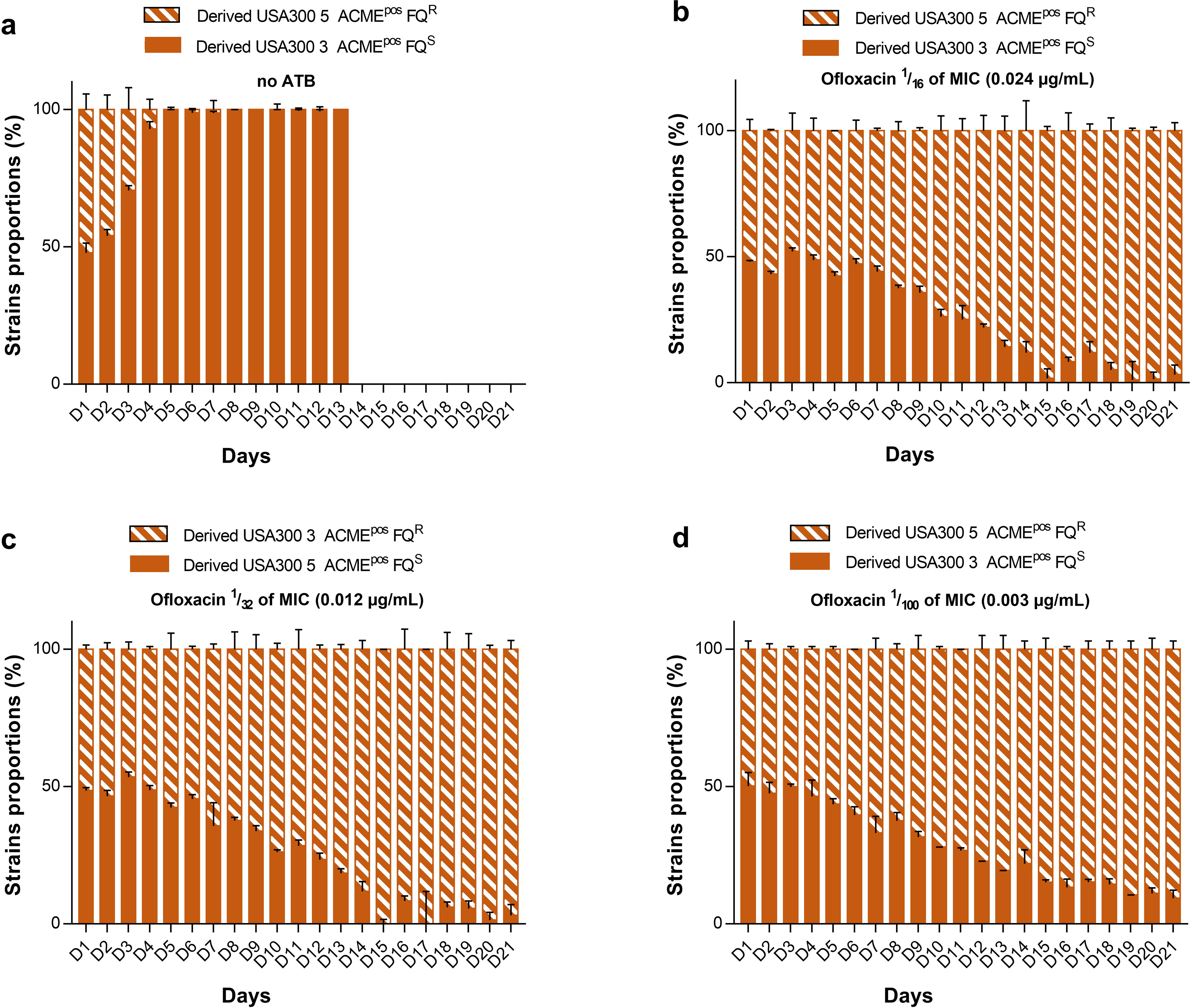
Potential impact of FQ resistance in competitive fitness of USA300. **(a)** Fluoroquinolone (FQ)-susceptible and -resistant strains of USA300, both ACME-positive, were co-cultivated for 21 days in BHI without antibiotics, or **(b)** containing ofloxacin at 1/16 of FQ MIC of the susceptible strain (0.024 μg/mL), **(c)** 1/32 MIC (0.012 μg/mL) or **(d)** 1/100 MIC (0.003 μg/ml) with daily subculture in fresh medium. The proportion of each strain was monitored daily with differential colony count based on selective agar inoculated with a calibrated amount of competitive mix. Competitive cultures were performed on three independent series (biological replicates), and each colony count or qPCR was repeated three times (technical replicates).

**Figure 7:**
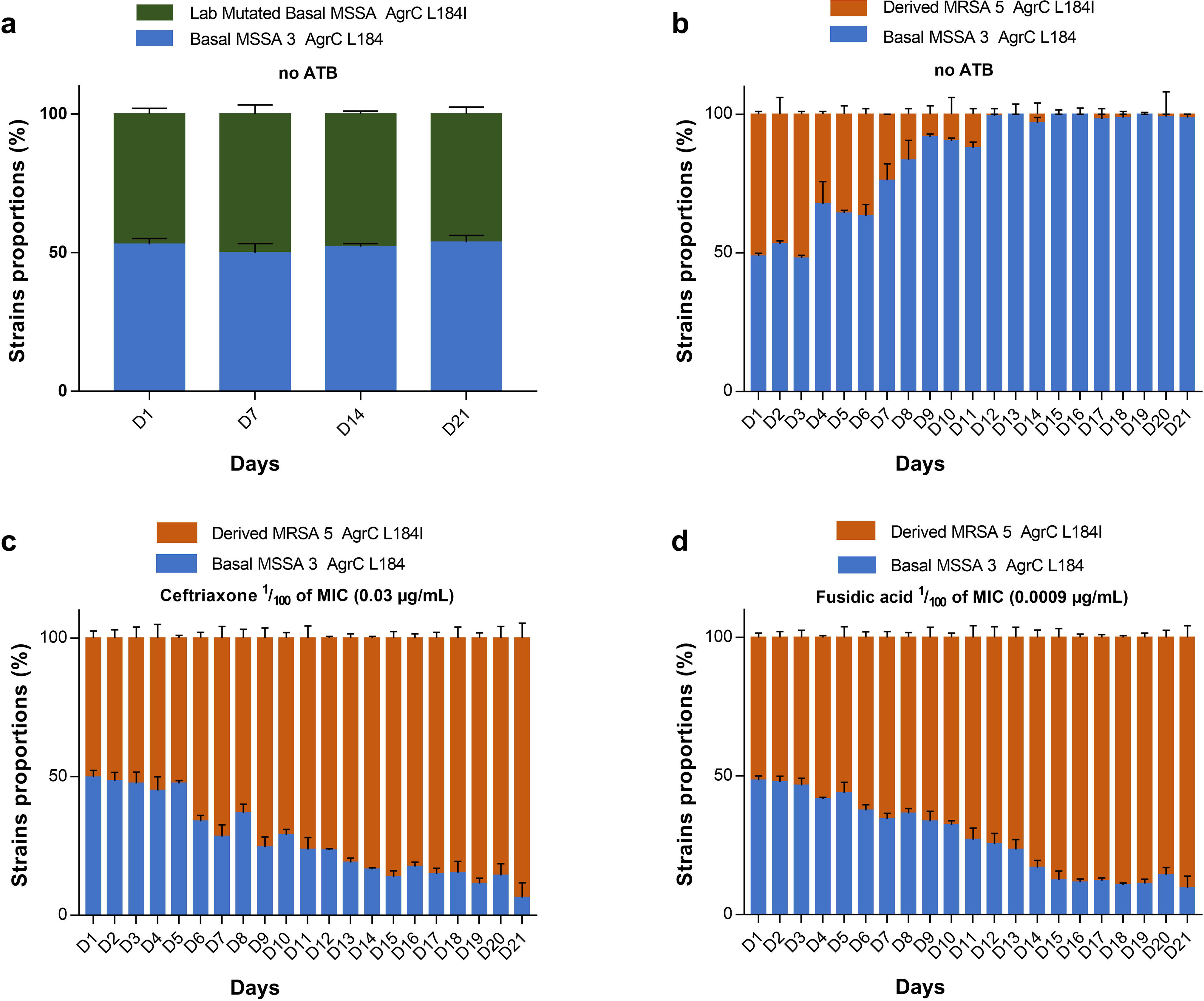
Potential impact of *mecA/fusB* acquisition and *agrC* mutation on competitive fitness of EU-ST80. **(a)** The *mecA*/*fus*B-negative (Basal MSSA 3) AgrC L184 was co-cultivated with its *agrC* derivative (Lab Mutated Basal MSSA) carrying the AgrC L184I mutation, **(b)** A *mecA*/*fus*B-negative (Basal MSSA 3) AgrC wild type (L184) strain and a *mecA*/*fus*B-positive (Derived MRSA 5) AgrC L84I strain were then co-cultivated for 21 days in BHI without antibiotics; same results were obtained with Basal MSSA 4 versus Derived MRSA 3 (data not shown), **(c) & (d)** The Basal MSSA 3 strain was co-cultivated with the Derived MRSA 5 strain for 21 days in BHI containing ceftriaxone or fusidic acid at 1/100 of MIC for the MSSA/*fus*B^neg^ strain (0.03 μg/mL or 0.0009 μg/ml_ respectively) with daily subculture in fresh medium. The proportion of each strain was monitored daily with differential colony count based on selective agar inoculated with a calibrated amount of competitive mix for **(b), (c), (d)** or by quantitative PCR due to the lack of discriminant antibiotic resistance marker for **(a).** The same results were obtained with antibiotics concentrations of 1/16 and 1/32 of their MICs and with the couple Basal MSSA 4 versus Derived MRSA 3 (data not shown). Competitive cultures were performed on three independent series (biological replicates), and each colony count or qPCR was repeated three times (technical replicates).

## DISCUSSION

Polyphyletic CA-MRSA emergence and spread at the end of the 20^th^ century (Vandenesch *et al.*, 2003), (Tristan *et al.*, 2007), remains a challenging issue. As pointed out by A-C. Uhlemann, “our understanding of how a clone [such as USA300 or EU-ST80] became established as an endemic pathogen within communities remains limited” (Uhlemann *et al.,* 2014). Increased expression of core-genome encoded virulence factors has been shown to be a common feature of CA-MRSA (Li *et al.*, 2010); we have investigated whether such characteristics varied along the longitudinal short-term evolution of CA-MRSA. However, by assessing transcription of virulence factors previously described as overexpressed among CA-MRSA lineages (Li *et al.*, 2010), we could detect only minor variation (ca. 1.5-fold increase) in virulence factor expression between USA300 strains when comparing ancestral and derived isolates of the North American clone (Fig. 3a); we cannot rule out however that, at the population level, these minor increases in virulence factor expression enhanced the success of the lineage, for instance by increasing cutaneous infection rate (the most common infections caused by *S.aureus*) and thus human-to-human transmission by skin contact as observed in prisons, sport team or men-having-sex-with-men (Planet *et al.*, 2015). Within this USA300 clone, 20 SNPs were identified for being under positive selection along successful evolution of this lineage (Glaser *et al.*, 2016); however, all the derived isolates assessed for doubling time carried these 20 SNPs. Therefore, despite being under positive selection they could not be reliable determinants of crude fitness evolution of the derived clade isolates as competitive fitness impairment observed along the phylogeny could not be explained by these genetic variations. Thus, the major variable feature of USA300 along the demography was the acquisition of ACME which is a now well-characterized mobile genetic element (MGE) acquired from *S. epidermidis* by horizontal gene transfer (Diep *et al.*, 2006), (Pi *et al.*, 2009), (Uhlemann *et al.*, 2014). Its multiple functions in resistance to acidic pH which enhances skin colonization, and as a factor promoting resistance to skin innate-immune defences (Thurlow *et al.*, 2012) makes it a very plausible contributor of the USA300 expansion (Planet, 2017). This is further strengthened by our findings of a shorter doubling-time of strains carrying ACME (Fig. 2a). In the case of EU-ST80, analysis of virulence factor expression along the demographical steps of the lineage showed that two derived isolates had a two-fold increase in *psm*α expression and another one had a 2.5-fold increase in PVL expression when compared to the ancestral isolates. As previously described (Stegger *et al.*, 2014), isolates of the basal and derived clades of this lineage were discriminated by four canonical SNPs. One was located in a non-coding region, two were synonymous SNPs, and one was a non-synonymous SNP located in *agrC*, the major virulence factor regulator involved in quorum sensing and biofilm production.

We focused our attention on this SNP located in the *agrC* gene, because of its potential association to fitness and colonization ability. This SNP resulted in a L184I amino acid change in the extracellular loop of the AgrC receptor (Stegger *et al.*, 2014) shared by all EU-ST80 isolates belonging to the derived clade. To investigate this point further, an ancestral ST80 (AgrC L184) was engineered by allelic replacement to carry the L184I substitution. Despite a slight increase of doubling time compared to its parental strain, the L184I change in AgrC did not translate into significant crude fitness variation (Mann-Whitney test, *P* = 0.343). Assessment of virulence factor expression revealed a two-fold increase of *psm*α in the Lab Mutated basal MSSA compared to its parental wild-type strain (Fig. 3b). AgrC L184I could therefore have a moderate impact on virulence. We further detected a strong and significant decrease in biofilm production associated with the AgrC L184I mutation (Fig. 4). Assessing which of these phenotypes (slight increase in PSMα or PVL, strong decrease in biofilm) was under selection remains speculative because they could be strongly dependent on the ecosystem in which selection has occurred. However, little is known regarding these ecological conditions since the current model for CA-MRSA ST80 lineage expansion places the acquisition of the AgrC L184I mutation in the early 1990s in strains originating from Sub-Saharan Western Africa, concomitantly with the acquisition of SCC*mec*IV and *fus*B (Stegger *et al.*, 2014). Alternatively, the AgrC L184I substitution might be a non-adaptive sequel – a genetic drift – parallel to the acquisition of SCC*mec*IV and *fus*B. Importantly, antibiotic resistances were associated with demographic expansion of EU-ST80 (acquisition of SCCmecIV and *fus*B) and also North American USA300 (acquisition of SCC *mec* and fluoroquinolone resistance) (Fig. 1) (Stegger *et al.*, 2014), (Glaser *et al.*, 2016). These resistance acquisitions were associated with a significant fitness cost as indicated by both extended doubling-time of the derived isolates (Fig. 2) and by the results of competition experiments where derived isolates were outcompeted by their basal counterparts (Fig. 5-7). These observations were in accordance with the classical fitness costs associated with de novo antibiotic resistance, specifically those selected at high antibiotic concentration (Martinez, 2009), (Andersson and Hughes, 2014). Conversely, they did not match the Bayesian evolutionary models of these lineages as strains belonging to the epidemic phase (derived clade) displayed the lowest *in vitro* competitive fitness, with each step of fitness decrease being associated with new acquisition of antibiotic resistance (Fig. 2, Table 1). However, the most striking observation was that extremely low concentrations of antibiotics (those for which resistance acquisition correspond to demographic expansion of the two lineages), totally reversed this fitness cost. Since both USA300 and EU-ST80 likely emerged in low income populations(Vandenesch *et al.*, 2003), (Martinez, 2009), (Planet, 2017), the role of antibiotic selective pressure was not initially considered to be the major trait under positive selection. However, increasing number of reports reveals the escalation of antibiotic as environmental pollutants originating from hospital wastewater, bulk drug producer wastewater and unused antibiotics dumped in landfills in countries without solid take-back programs (Naimi *et al.*, 2003), (Thurlow *et al.*, 2012), (Larsson, 2014), (Gothwal and Shashidhar, 2015), (See *et al.*, 2017). From these sources, in which antibiotics such as fluoroquinolones can reach concentrations ranging from 3 ng/L to 240 μg/L (Van Doorslaer *et al.*, 2014), antibiotics are disseminated in various environmental matrices such as surface water, soil, sediments, and eventually living organism including livestock (Van Doorslaer *et al.*, 2014). Hence, community settings, even in remote populations, can be exposed to low-level concentrations of various antibiotics that could have promoted the expansion of CA-MRSA at least by enriching for resistant bacteria (Andersson and Hughes, 2014), if not selecting for de novo resistance, the latter being typically associated with no fitness cost (Gullberg *et al.*, 2011), (Andersson and Hughes, 2014), (Westhoff *et al.*, 2017). Here, we demonstrate with competition experiments that the biological cost of antibiotic resistance (to beta-lactams, fusidic-acid and fluoroquinolone) is entirely reversed in the presence of trace amounts of antibiotics. Previous studies based on multidrug resistant plasmids showed that, for specific combinations of drugs, each new compound added, lowered the minimal selective concentration of the others (Gullberg *et al.,* 2014). However antibiotic resistance acquisitions (both by horizontal transfer of resistance genes and by mutations) are the genetic events that best match the variation of the demography in both lineages (Fig. 1a & b). Altogether, our findings support a model of antibiotic use, misuse and pollution as a major driving force for the emergence and expansion of CA-MRSA. In conclusion, CA-MRSA dynamics appear to be ruled by a complex interplay between resistance, virulence and fitness cost in which the contribution of anthropogenic activities is substantial.

## ACKNOWLEDGEMENTS

We thank Alex Van Belkum for fruitful discussion, and the technicians and engineers of the French National Reference Center for Staphylococci for their skilful contribution. This work was not supported by specific grants. The salaries (C-A. G., A. T., P. M-S., Y. B., M. B., F. L., F. V.) were supported by the University of Lyon, *Hôpitaux de Lyon* and by *Santé Publique France* under the funding of the French National Reference Center for Staphylococci. The funders had no role in study design, data collection and interpretation, or the decision to submit the work for publication.

## CONFLICT OF INTEREST

The authors declare no conflict of interest.

Supplementary information is available at the ISME Journal’s website

**<u>Table SI:</u> Strains pairs used for competitive fitness study**

Strains pairs listed according to their lineage assignment, with discriminant parameters (“Striking difference”).

**<u>Table S2:</u> Primers used for qPCR and RT-qPCR**

Primers list including target genes used for resistance-based strain discrimination (*mecA, tetK, grlA*), phylogenic clade discrimination (*agrC*, *arc*A-ACME), virulence factor expression assay (*RNAIII*, *hla*, *hlgC*, *lukS-PV*, *PSMα*), engineering of the Lab Mutated basal MSSA (*agrC*2912, 555, 4238, 544), and standardization *igyr*)

## REFERENCES

Andersson DI, Hughes D. (2014). Microbiological effects of sublethal levels of antibiotics. Nat Rev Microbiol 12: 465–478.

Arnaud M, Chastanet A, Débarbouillé M. (2004). New Vector for Efficient Allelic Replacement in Naturally Nontransformable, Low-GC-Content, Gram-Positive Bacteria. Appl Environ Microbiol 70:6887–6891.

Chambers HF. (2001). The changing epidemiology of Staphylococcus aureus? Emerg Infect Dis 7: 178–182.

Chavez TT, Decker CF. (2008). Health Care-Associated MRSA Versus Community-Associated MRSA. Dis Mon 54: 763–768.

David MZ, Daum RS. (2010). Community-Associated Methicillin-Resistant Staphylococcus aureus: Epidemiology and Clinical Consequences of an Emerging Epidemic. Clin Microbiol Rev 23: 616–687.

Diep BA, Gill SR, Chang RF, Phan TH, Chen JH, Davidson MG, et al. (2006). Complete genome sequence of USA300, an epidemic clone of community-acquired meticillin-resistant Staphylococcus aureus. The Lancet 367: 731–739.

Glaser P, Martins-Simões P, Villain A, Barbier M, Tristan A, Bouchier C, et al. (2016). Demography and Intercontinental Spread of the USA300 Community-Acquired Methicillin-Resistant Staphylococcus aureus Lineage. mBio 7. e-pub ahead of print, doi: 10.1128/m Bio.02183-15.

Gothwal R, Shashidhar T. (2015). Antibiotic Pollution in the Environment: A Review. CLEAN – Soil Air Water 43:479–489.

Gullberg E, Albrecht LM, Karlsson C, Sandegren L, Andersson DI. (2014). Selection of a Multidrug Resistance Plasmid by Sublethal Levels of Antibiotics and Heavy Metals. mBio 5: e01918–14.

Gullberg E, Cao S, Berg OG, llback C, Sandegren L, Hughes D, et al. (2011). Selection of Resistant Bacteria at Very Low Antibiotic Concentrations. PLOS Pathog 7: el002158.

Horsburgh MJ, Aish JL, White IJ, Shaw L, Lithgow JK, Foster SJ. (2002). σB Modulates Virulence Determinant Expression and Stress Resistance: Characterization of a Functional rsbU Strain Derived from Staphylococcus aureus 8325-4. J Bacteriol 184: 5457–5467.

Larsson DGJ. (2014). Antibiotics in the environment. Ups J Med Sci 119:108–112.

Li M, Cheung GYC, Hu J, Wang D, Joo H-S, DeLeo FR, et al. (2010). Comparative Analysis of Virulence and Toxin Expression of Global Community-Associated Methicillin-Resistant Staphylococcus aureus Strains. J Infect Dis 202:1866–1876.

Lowy FD. (1998). Staphylococcus aureus Infections. N Engl J Med 339: 520–532.

Ma XX, Ito T, Tiensasitorn C, Jamklang M, Chongtrakool P, Boyle-Vavra S, et al. (2002). Novel Type of Staphylococcal Cassette Chromosome mec Identified in Community-Acquired Methicillin-Resistant Staphylococcus aureus Strains. Antimicrob Agents Chemother 46: 1147–1152.

Martinez JL. (2009). The role of natural environments in the evolution of resistance traits in pathogenic bacteria. Proc R Soc B Biol Sci 276: 2521–2530.

Mediavilla JR, Chen L, Mathema B, Kreiswirth BN. (2012). Global epidemiology of community-associated methicillin resistant Staphylococcus aureus (CA-MRSA). Curr Opin Microbiol 15: 588–595.

Naimi TS, LeDell KH, Como-Sabetti K, Borchardt SM, Boxrud DJ, Etienne J, et al. (2003). Comparison of Community- and Health Care-Associated Methicillin-Resistant Staphylococcus aureus Infection. JAMA 290: 2976–2984.

Okuma K, Iwakawa K, Turnidge JD, Grubb WB, Bell JM, O’Brien FG, et al. (2002). Dissemination of New Methicillin-Resistant Staphylococcus aureus Clones in the Community. J Clin Microbiol 40: 4289–4294.

O’Neill A j. (2010). Staphylococcus aureus SH1000 and 8325-4: comparative genome sequences of key laboratory strains in staphylococcal research. LettAppl Microbiol 51: 358–361.

Pi B, Yu M, Chen Y, Yu Y, Li L. (2009). Distribution of the ACME-arcA gene among meticillin-resistant Staphylococcus haemolyticus and identification of a novel ccr allotype in ACME-arcA-positive isolates. J Med Microbiol 58: 731–736.

Planet PJ. (2017).Life After USA300: The Rise and Fall of a Superbug. J Infect Dis 215: S71–S77.

Planet PJ, Diaz L, Kolokotronis S-O, Narechania A, Reyes J, Xing G, et al. (2015). Parallel Epidemics of Community-Associated Methicillin-Resistant Staphylococcus aureus USA300 Infection in North and South America. J Infect Dis 212:1874–1882.

Regev-Yochay G, Trzciński K, Thompson CM, Malley R, Lipsitch M. (2006). Interference between Streptococcus pneumoniae and Staphylococcus aureus: In Vitro Hydrogen Peroxide-Mediated Killing by Streptococcus pneumoniae. J Bacteriol 188: 4996–5001.

Reynolds J, Wigneshweraraj S. (2011). Molecular insights into the control of transcription initiation at the Staphylococcus aureus agr operon. J Mol Biol 412: 862–881.

Rosenstein R, Nerz C, Biswas L, Resch A, Raddatz G, Schuster SC, et al. (2009). Genome Analysis of the Meat Starter Culture Bacterium Staphylococcus carnosus TM300. Appl Environ Microbiol 75: 811–822.

See I, Wesson P, Gualandi N, Dumyati G, Harrison LH, Lesher L, et al. (2017). Socioeconomic Factors Explain Racial Disparities in Invasive Community-Associated Methicillin-Resistant Staphylococcus aureus Disease Rates. Clin Infect Dis 64: 597–604.

Stegger M, Wirth T, Andersen PS, Skov RL, De Grassi A, Simões PM, et al. (2014). Origin and Evolution of European Community-Acquired Methicillin-Resistant Staphylococcus aureus. mBio 5. e-pub ahead of print, doi: 10.1128/mBio.01044-14.

Thurlow LR, Joshi GS, Clark J R, Spontak JS, Neely CJ, Maile R, et al. (2013). Functional Modularity of the Arginine Catabolic Mobile Element Contributes to the Success of USA300 Methicillin-Resistant Staphylococcus aureus. Cell Host Microbe 13:100–107.

Thurlow LR, Joshi GS, Richardson AR. (2012). Virulence Strategies of the Dominant USA300 Lineage of Community Associated Methicillin Resistant Staphylococcus aureus (CA-MRSA). FEMS Immunol Med Microbiol 65: 5–22.

Tristan A, Bes M, Meugnier H, Lina G, Bozdogan B, Courvalin P, et al. (2007). Global Distribution of Panton-Valentine Leukocidin-positive Methicillin-resistant Staphylococcus aureus, 2006. Emerg Infect Dis 13: 594–600.

Uhlemann A-C, Dordel J, Knox JR, Raven KE, Parkhill J, Holden MTG, et al. (2014). Molecular tracing of the emergence, diversification, and transmission of S. aureus sequence type 8 in a New York community. Proc Natl Acad Sci USA 111: 6738–6743.

Van Doorslaer X, Dewulf J, Van Langenhove H, Demeestere K. (2014). Fluoroquinolone antibiotics: An emerging class of environmental micropollutants. Sci Total Environ 500–501: 250–269.

Vandenesch F, Naimi T, Enright MC, Lina G, Nimmo GR, Heffernan H, et al. (2003). Community-Acquired Methicillin-Resistant Staphylococcus aureus Carrying Panton-Valentine Leukocidin Genes: Worldwide Emergence. Emerg Infect Dis 9: 978–984.

Westhoff S, van Leeuwe TM, Qachach O, Zhang Z, van Wezel GP, Rozen DE. (2017). The evolution of no-cost resistance at sub-MIC concentrations of streptomycin in Streptomyces coelicolor. ISMEJ 11:1168–1178.

Zhang Y-Q, Ren S-X, Li H-L, Wang Y-X, Fu G, Yang J, et al. (2003). Genome-based analysis of virulence genes in a non-biofilm-forming Staphylococcus epidermidis strain (ATCC 12228). Mol Microbiol 49: 1577–1593.

